# Allee effects mediate the impact of land-use change on the thermal niche of social species

**DOI:** 10.1101/2022.10.11.511768

**Authors:** Shih-Fan Chan, Dustin R. Rubenstein, Tsung-Wei Wang, Ying-Yu Chen, I-Ching Chen, Dong-Zheng Ni, Wei-Kai Shih, Sheng-Feng Shen

## Abstract

Land-use change not only affects habitat availability, it can also reduce population density and limit opportunities for interactions with conspecifics, further influencing species resilience to environmental challenges. For social species whose conspecific interactions are typically cooperative in nature, little is known about how land-use change influences demography and social behavior, and how this interaction impacts a species’ climatic niche. Here, we develop a spatially explicit, individual-based model to explore how land-use changes influence population size and niche width in social organisms through the Allee effect, the positive impact of higher population density on individual fitness. We then empirically test key model predictions by studying the distribution and cooperative behavior of burying beetles (*Nicrophorus nepalensis*) along elevational gradients in Taiwan. In support of our model predictions, we find that beetle densities are lower in areas of greater land-use change, making it harder for individuals in these hotter environments to form cooperative groups to compete against blowflies, their primary interspecific competitor. Consequently, the beetles’ lower distributional boundary is higher in areas with greater land-use change, indicating that the beetles’ thermal niche is reduced via Allee effects in human-altered landscapes. Ultimately, land-use change not only reduces habitat availability, it also shrinks the thermal niche of social species, making them more vulnerable to climate change.

## Introduction

Ongoing global climate change is causing dramatic biodiversity loss and species redistribution across the globe (Parmesan & Yohe 2003; Chen *et al*. 2011; Pecl *et al*. 2017). The direct effects of climate change, including declines in the local population abundance, local population extinction, and large-scale polar or upward shifts in distributional ranges have been widely explored (Parmesan 2006). However, an increasing number of studies have focused on the interactions between climate and other biodiversity stressors (Brook *et al*. 2008), particularly anthropogenic land-use change, which also extensively reshapes global species distributions and ecosystem functioning (Foley *et al*. 2005; Haddad *et al*. 2015; Newbold *et al*. 2015). Accumulating evidence suggests that land-use change reinforces the negative impacts of climate change, resulting in deleterious impacts greater than the sum of the individual contributions of these two threatening processes (Oliver & Morecroft 2014; Oliver *et al*. 2015; Guo *et al*. 2018; Peters *et al*. 2019; Elsen *et al*. 2020). Yet, the mechanism by which this interactive process works remains largely unexplored (Schulte to Bühne *et al*. 2021).

Critical to understanding how climate and land-use change synergistically affect organismal population abundance is to determine how heterogenous habitat loss affects organisms that live in the remaining pristine habitats patches (Travis 2003; Hof *et al*. 2011; Oliver & Morecroft 2014). Most theories explaining how climate and land-use change synergistically affect organismal population abundance have been largely based on the assumption that an organism’s climatic niche is not affected by land-use change. The idea for this comes from Hutchinson’s distinction between niche and biotope (Hutchinson 1957; Colwell Robert & Rangel Thiago 2009). The niche represents the n-dimension environmental attributes (e.g., temperature and humidity) required for a species’ positive population growth rate. In contrast, the biotope represents physical space, such as the habitat in which organisms are distributed (Colwell Robert & Rangel Thiago 2009). In other words, land-use change intuitively reduces habitat (biotope) availability (Platts *et al*. 2019) and connectivity (Hof *et al*. 2011; Senior *et al*. 2019), and prevents organisms from tracking suitable climates (niche) to keep pace with temperature change (Travis 2003; Tucker *et al*. 2018). However, an alternative but rarely considered possibility is that land-use change, after reducing available habitat (biotope) and causing an initial reduction in population density, then causes negative effects on social behaviors that are necessary for individual survival and reproduction, including mate searching (Wells *et al*. 1998), group foraging (Grünbaum & Veit 2003), or cooperation (Courchamp *et al*. 1999).

This inverse density-dependent relationship in which smaller population densities are associated with lower average individual fitness is often referred to as the Allee effect (Allee 1927; Allee 1931; Courchamp *et al*. 1999; Stephens & Sutherland 1999), a hallmark of social species that form groups. The Allee effect of reducing population density, driven by land-use change, may also reduce ecological niche width by making it more difficult for organisms to find mates or to cooperate against adverse conditions, among other mechanisms. Consequently, land-use change may not only directly impact organism by reducing the amount of suitable available habitats, but also by reducing the fitness of organisms living in the remaining pristine habitats in the local area. Yet, to our knowledge, no study has considered the possibility that in addition to affecting population density, reducing biotope may also affect niche space through Allee effects. Given that Allee effects are a common phenomenon in nature and occur in a diversity of vertebrate and invertebrate animal species (Courchamp *et al*. 1999; Stephens & Sutherland 1999; Angulo *et al*. 2007; Angulo *et al*. 2018), it is likely that the impact of land-use change on organisms under climate change has therefore been underestimated.

Here, we first construct a spatially explicit, individual-based model to explore how land-use change affects population density, which then further impacts the thermal niche in terms of distribution, population density, and breeding performance, of social organisms through the Allee effect. We then test the model’s key predictions by examining the elevational distribution and breeding performance of the Asian burying beetle (*Nicrophorus nepalensis*) across two elevational gradients in Taiwan that vary in their degree of land-use change and the amount of unaltered pristine forest. Because burying beetles use vertebrate carcasses as the sole resource for reproduction (Scott 1998), they represent valuable “bonanza resources” for many *Necrophagous* species (Wilson 1975). The reproductive success of *N. nepalensis* is strongly limited by interspecific competition for these resources, especially from blowflies (Scott 1994; Sun *et al*. 2014; Chan *et al*. 2019; Chen *et al*. 2020). *N. nepalensis* overcomes the challenge of interspecific competition with blowflies through intraspecific cooperation, which allows the beetles to successfully reproduce in warmer environments at lower elevations characterized by stronger blowfly competition (Sun *et al*. 2014; Liu *et al*. 2020; Tsai *et al*. 2020a). Intraspecific cooperation thus helps burying beetles to expand their thermal niche from cooler environments at higher elevations where blowflies are scarce or absent to warmer environments at lower elevations where blowflies are more abundant. Furthermore, the reproductive success of *N. nepalensis* is also limited by its ability to locate carcasses, especially in colder environments at higher elevations where extended movement is costly (Chan *et al*. 2019; Liu *et al*. 2020). Although we study a social species here because intraspecific cooperation and group formation are the primary causes of Allee effect, it is important to emphasize that Allee effect can be applied to nonsocial species as long as the population density has a positive effect on population growth (Courchamp *et al*. 1999; Stephens & Sutherland 1999; Angulo *et al*. 2007; Angulo *et al*. 2018).

Specifically, we use theoretical models and empirical studies to test (i) the Allee effect hypothesis, which states that in a pristine landscape with limited land-use change, social species will have higher population densities and be more likely to cooperate with other individuals to cope with environmental challenges such as harsh conditions or interspecific competition. Therefore, we predict that social species will have larger group sizes in pristine landscapes than in human-altered landscapes when faced with environmental challenges. As a result, social species can adapt to a greater range of environmental conditions and will therefore have a greater geographic range and a wider niche width. Alternatively, under (ii) the fixed niche width hypothesis, land-use change will only affect habitat availability, and therefore a species’ niche width will only be affected by the physiological and behavioral characteristics of that species, not by its population density. Therefore, we predict that although population densities will be higher in pristine than human-altered landscapes, species geographic range sizes and niche widths will be similar in both landscapes. Investigating the effects of land-use change-driven population density declines on the reproduction and distribution of burying beetles along elevation (and thermal) gradients provides an ideal opportunity to determine how demographic changes at the landscape scale alter species responses to climate change through the Allee effect. In addition, our study provides a set of quantifiable metrics that will facilitate the development of large-scale conservation strategies for habitat management in the face of climate change.

## Results

### Theoretical model

We begin by developing an individual-based model to examine the effects of land-use change on population density and the climatic niche width of social organisms (Fig. 1). We assume that breeding performance is determined by the thermal performance curve (TPC) when individuals reproduce solitarily (Figure 1— Figure Supplement 1A). We further assume that cooperation enables individuals to improve their breeding performance, and because burying beetles experience greater interspecific competition from flies at higher temperatures (Sun *et al*. 2014; Liu *et al*. 2020), that the benefit of cooperation is greater at higher temperatures (Figure 1— Figure Supplement 1B). However, as the number of group members grows too large, the cooperative benefits will eventually saturate and stop increasing (Figure 1— Figure Supplement 1B). Therefore, our model explores the relationship between the physiological performance of individuals with the cooperative benefits at different environmental temperatures. We can see that when the environmental temperature exceeds an individual’s physiologically optimal temperature, (i) the breeding performance of individuals in a social group remains poor (Figure 1— Figure Supplement 1C), even when the cooperative benefits are high (Figure 1— Figure Supplement 1B), and (ii) group productivity is higher in larger social groups (Figure 1— Figure Supplement 1D).

**Figure 1.**
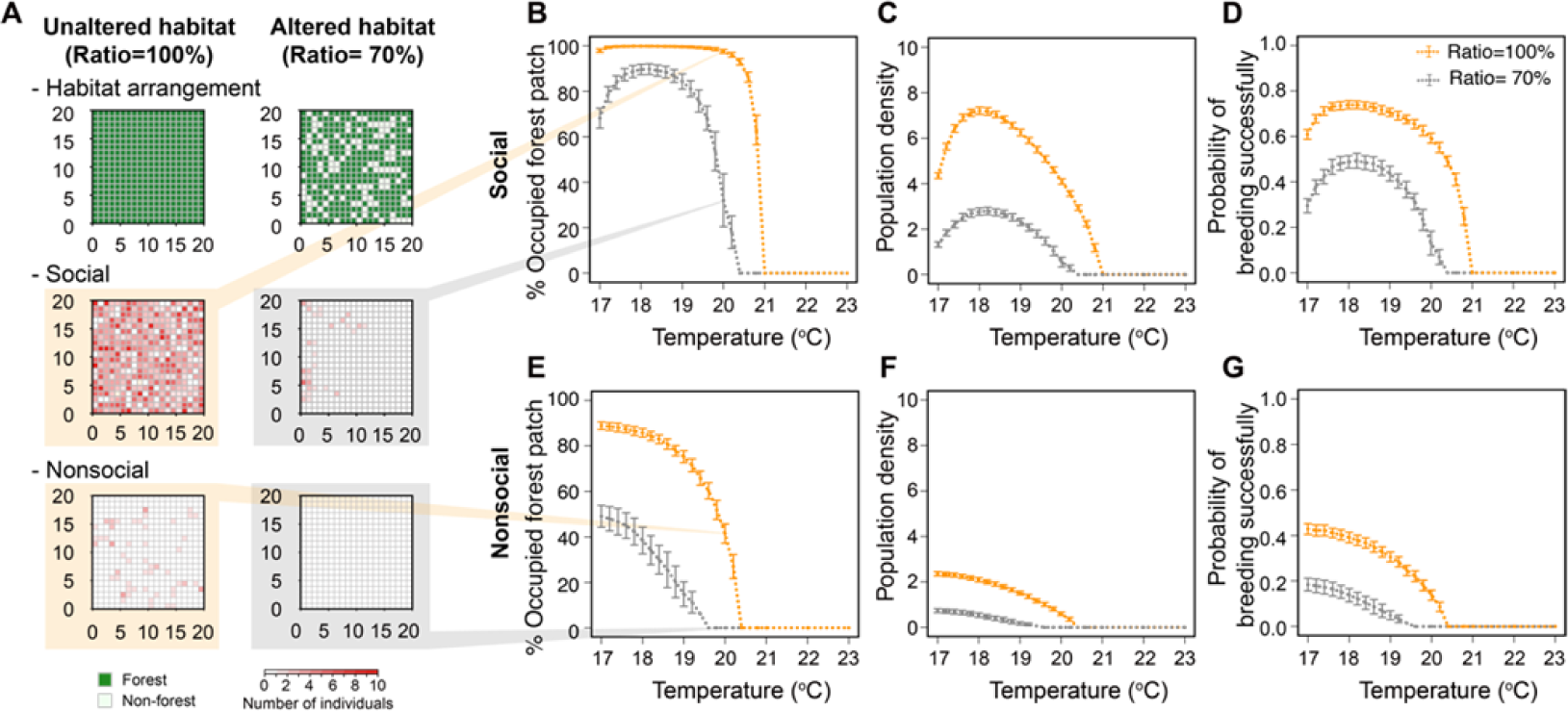
Individual-based modeling for the impacts of land-use change on the thermal niche of social and nonsocial populations. (A) A pair of examples of different forest habitats (with a different habitat ratio) for simulations, with a set of simulated results at the temperature of 20°C. (B-D) Simulation results for patch occupation rate (B), population density (C), and probability of breeding successfully (D) along the temperature gradient in continuous and fragmented habitats in social species. (E-G) Simulation results for patch occupation rate (E), population density (F), and probability of breeding successfully (G) along the temperature gradient in continuous and fragmented habitats in nonsocial species. In (A), each grid represents a habitat patch. In (B-G), points and error bars represent means and standard deviations obtained from 200 simulation runs.

For simplicity, we begin with a population that is composed entirely of cooperative individuals. For comparison, however, we also constructed a model with the same settings, but with a population composed entirely of noncooperators. Because the impacts of land-use change that we want to investigate occur at ecological time scales, we do not consider the possibility that cooperative behavior can evolve. Instead, we assume that the degree of individual cooperation is fixed. Thus, the productivity of a group is mainly influenced by the group size within a patch and environmental temperature. More individuals produce more group resources until the size of the group is larger than the optimal size. We assume that the environmental resources are finite, and therefore, cooperative groups cannot produce more group resources indefinitely.

The effect of environmental temperature on an individual’s reproductive success in a non-cooperative situation is determined by the thermal performance curve (TPC). For simplicity, we only explore the warm distribution boundary (i.e., low elevation) with this model, though we note that the results would be qualitatively similar if we simulate both the warm and cold (i.e., high elevation) distribution boundaries together. Thus, we only simulate a situation in which the ambient temperature is higher than the optimum temperature of the species, adopting the thermal performance curve as:

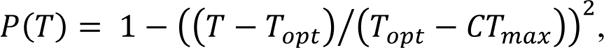

where T is the environmental temperature, T_opt_, is the species’ optimum temperature, and CT_max_ is the critical thermal maximum (Deutsch *et al*. 2008; Vasseur *et al*. 2014). We set T_opt_ to 16.7 °C and CT_max_ to 23.0 °C, based on the published data of *N. nepalensis* (Tsai *et al*. 2020a). We simulate the ambient temperature range of 17-23°C at 0.2°C intervals. Although our model is parametrized based on the biology of *N. nepalensis*, we emphasize that it is applicable to any species (social or otherwise) whose temperature-dependent fitness components can be described by TPCs.

For each temperature condition, we set a spatial range of 400 patches and randomly select the suitable forest habitats according to the habitat ratio (e.g., a habitat ratio of 0.7 means that 280 patches are randomly selected as suitable forest habitats for breeding) (Fig. 1A). For each simulation, the starting population size is set to 0.6 times the number of patches (i.e., 240 individuals) and is randomly distributed throughout the space. At each time step of the simulation, cooperating individuals in the same patch produce group resources and reproduce when they occur in suitable habitats. However, in unsuitable habitats, individuals simply die and cannot reproduce. For simplicity, we consider asexual reproduction. After an individual produces offspring, the offspring will randomly disperse into a new patch. The total time step of the simulation is 2,000 to ensure that the system reaches a steady state. We record the final number of patches with individuals, the final population size and population density (i.e., the final population size/the number of patches suitable for reproduction), and the number of offspring produced by an individual in each patch. We also calculate the per capita offspring production accordingly. We repeat the simulation 200 times for each environmental condition and calculate the mean and standard deviation of the 200 results.

We found that the temperature limit of the distribution of cooperative populations in human-altered forests (forest habitat availability = 70%) was approximately 0.6 degrees lower than that of pristine forest habitats (forest habitat availability = 100%), in terms of the number of occupied patches (Fig. 1B), population density (Fig. 1C) and of breeding performance of social groups (Fig. 1D). We also found that beetles had larger group sizes in pristine forest habitat (Figure 1— Figure Supplement 2), suggesting that land-use changes reduce the thermal niche and the distribution ranges of the species because larger population sizes facilitate cooperation of social populations in pristine forest habitats, which in turn positively affects reproduction (i.e., an Allee effect) (Figure 1— Figure Supplement 3). When we simulated a non-cooperative population for comparison, we found that stochastic extinctions of small, nonsocial populations can also cause Allee effects (Fig. 1E-G). However, the positive effect of greater population density is smaller in nonsocial populations than social ones. Thus, under the same environmental conditions, the thermal niche width of nonsocial populations was narrower than the number of occupied patches (Fig. 1E), the population density (Fig. 1F), and the breeding performance (Fig. 1G) of social populations. Together, our simulations suggest that land-use change significantly reduces the thermal niche width of species through the Allee effect.

### Empirical results

Next, we tested the key predictions of our model by studying *N. nepalensis* along elevational gradients in Taiwan. Based on the model results, we predict that in areas with more land-use change, the density of burying beetles will decrease, making it more difficult for them to form cooperative groups. Thus, the beetles’ thermal niche will shrink to the point that beetles can only reproduce successfully in cooler environments where competition with blowflies is lower. For this reason, the lower boundary of the beetle’s elevation distribution will occur at higher elevations in areas with greater land-use change. Conversely, if land-use change only affects the availability of suitable habitats (and not population density or the propensity to cooperate), we predict that the thermal niches of burying beetles will not be affected by land-use change.

We first quantified forest cover along two mountain gradients in Mt. Hehuan, Taiwan (Fig 2A). We found that the forest on the western slope of Mt. Hehuan has been affected by extensive human cultivation and forest fragmentation. In contrast, the eastern slope of Mt. Hehuan is located within Taroko National Park and is covered by well-protected (pristine) natural forests, resulting in significantly lower forest cover below 2500 m on the western slope than on the eastern slope (Linear model, interaction Region*Elevation, Region*Elevation^2^, and main effect of Region, all P<0.001, n=71; Fig. 2B; Table S1). However, we found that the relationship between temperature and elevation under the remaining forests was similar on both slopes, indicating that the difference in forest cover only slightly affected the average daily temperature under the canopy on either slope (LMM, interaction Region*Elevation, P=0.09; main effect of Region, P=0.29, n=4849; Fig. 2C; Table S2).

**Figure 2.**
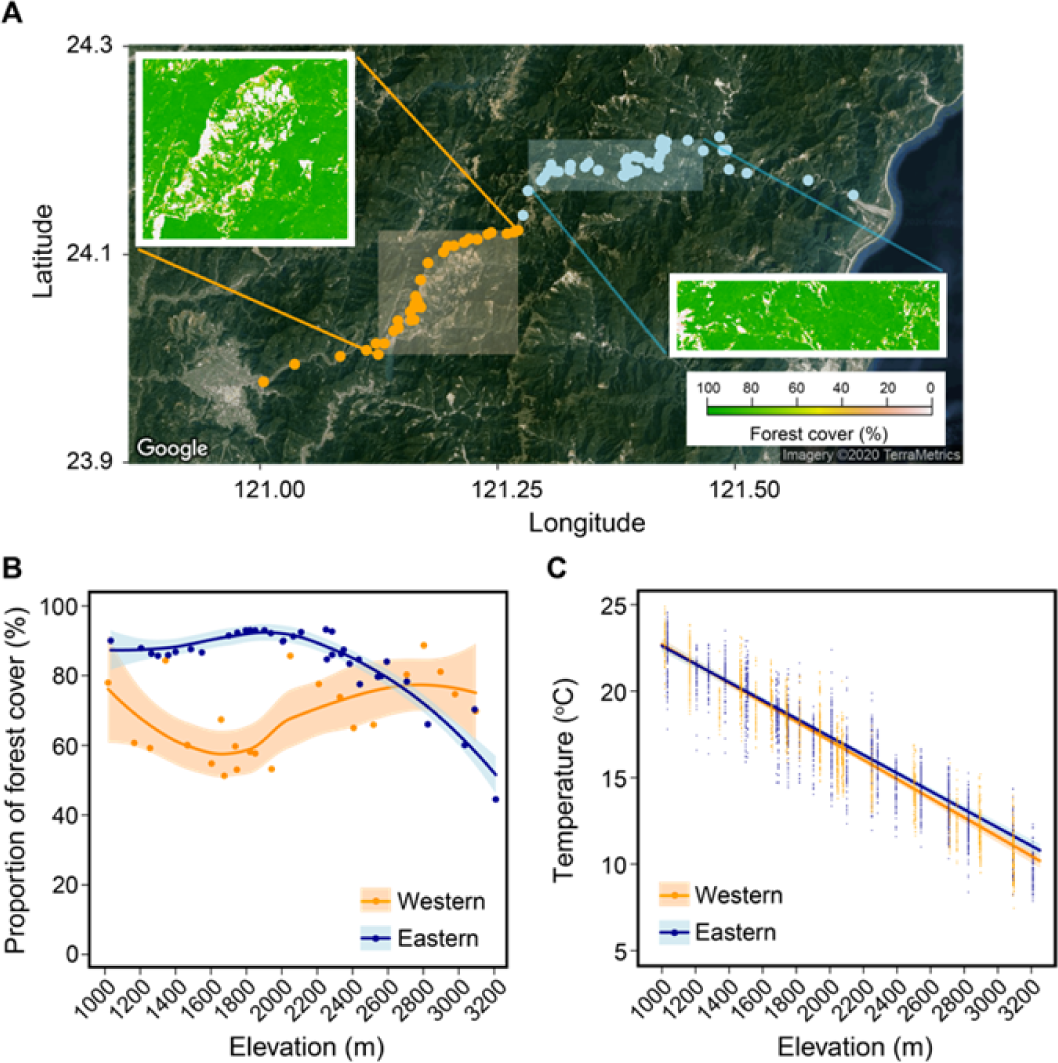
Environmental features in the study region. (A) Satellite image for the study region obtained from Google Maps. Orange and light blue points represent the sampling sites along the western and eastern slopes of Mt. Hehuan, respectively. Semitransparent rectangle areas represent the regions in which we carried out breeding experiments. The insets indicate the distribution of forest cover in the semitransparent rectangle areas. (B) The difference in elevational trends in the percentage of forest cover between the two slopes. (C) Elevational trends in environmental temperature along the two slopes. Trends and 95% confidence intervals (shaded areas) are estimated from LOWESS (in B) and GLMM (in C).

Next, we wanted to investigate the effects of land-use change on the density and breeding performance of burying beetles to understand possible Allee effects on their breeding thermal niche. We found that the higher the forest cover, the greater the beetle density (GLMM, Forest cover, P<0.001, n=372; Fig. 3A; Table S3A). Although burying beetle densities were lower on western slopes (with greater land-use change) than on eastern slopes (with lower land-use change) at lower elevations with higher temperatures, there were little differences in densities between the eastern and western slopes at higher elevations with lower temperatures (GLMM, interaction Temperature*Region, P=0.09; main effect of Region, P<0.01, n=372; Fig. 3B; Table S3B). These results suggest that land-use change in warmer environments at lower elevations has a greater negative impact on burrowing beetle densities than in colder environments at higher elevations.

**Figure 3.**
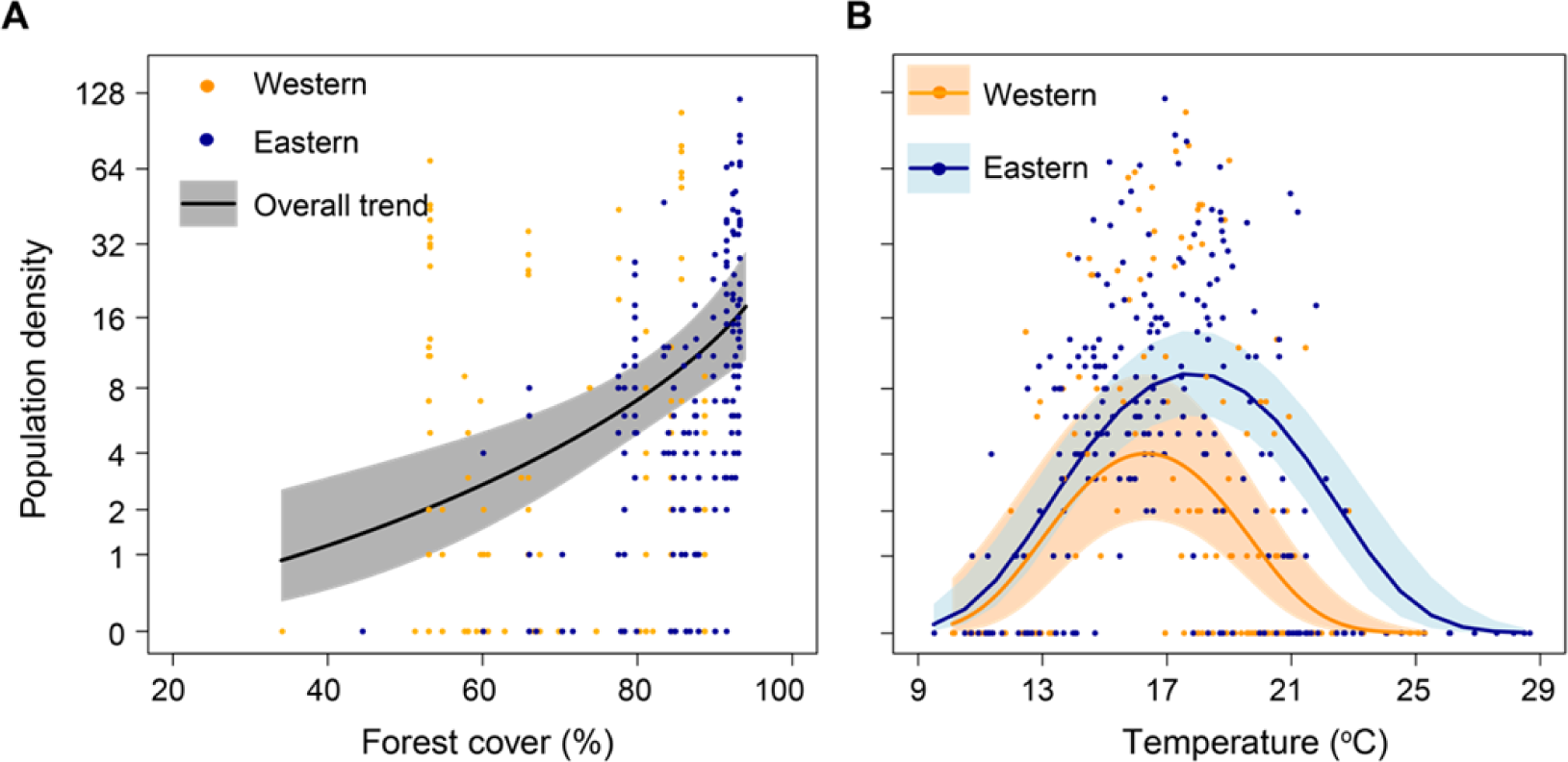
Factors affecting population density of burying beetles. (A) The relationship between forest cover and beetle density. (B) The relationship between temperature and beetle population density. All trends and 95% confidence intervals (shaded areas) are estimated from GLMM.

We then conducted a set of field breeding experiments along the two elevational gradients to investigate how differences in population density caused by land-use changes affect the thermal niche of burying beetles. We found that the higher beetle population density driven by a greater percentage of forest cover further affected beetle breeding performance at different temperatures (i.e., breeding thermal niche). That is, the reproductive success of burying beetles increased with increasing population density (GLMM, Population density, P<0.001, n=322; Fig. 4A; Table S4). In other words, high density burying beetle populations had a higher probability of breeding successfully than low density populations at the same temperature (Fig. 4B; Table S4). We further analyzed the behavioral mechanism of how population density affects the reproductive success of burying beetles and found that higher population densities resulted in larger cooperative groups than populations with lower densities at low elevations with higher temperatures (GLMM, interaction Population density*Temperature, P<0.01; main effect of Population density, P<0.001, n=199; Fig. 4C; Table S5). In addition, higher beetle population densities also resulted in earlier carcass discoveries by the beetles (GLMM, interaction Population density*Temperature, P=0.07; main effect of Population density, P<0.001, n=210; Fig. 4D; Table S6). Together, we used piecewise SEM analysis (Shipley 2009; Lefcheck 2016) to show that these two behavioral mechanisms increased the beetles’ reproductive success (Fig. 4E; Table S7). Therefore, we found that deforestation reduces the population density of burying beetles, which in turn shrinks the size of the beetles’ breeding thermal niche space through the Allee effect.

**Figure 4.**
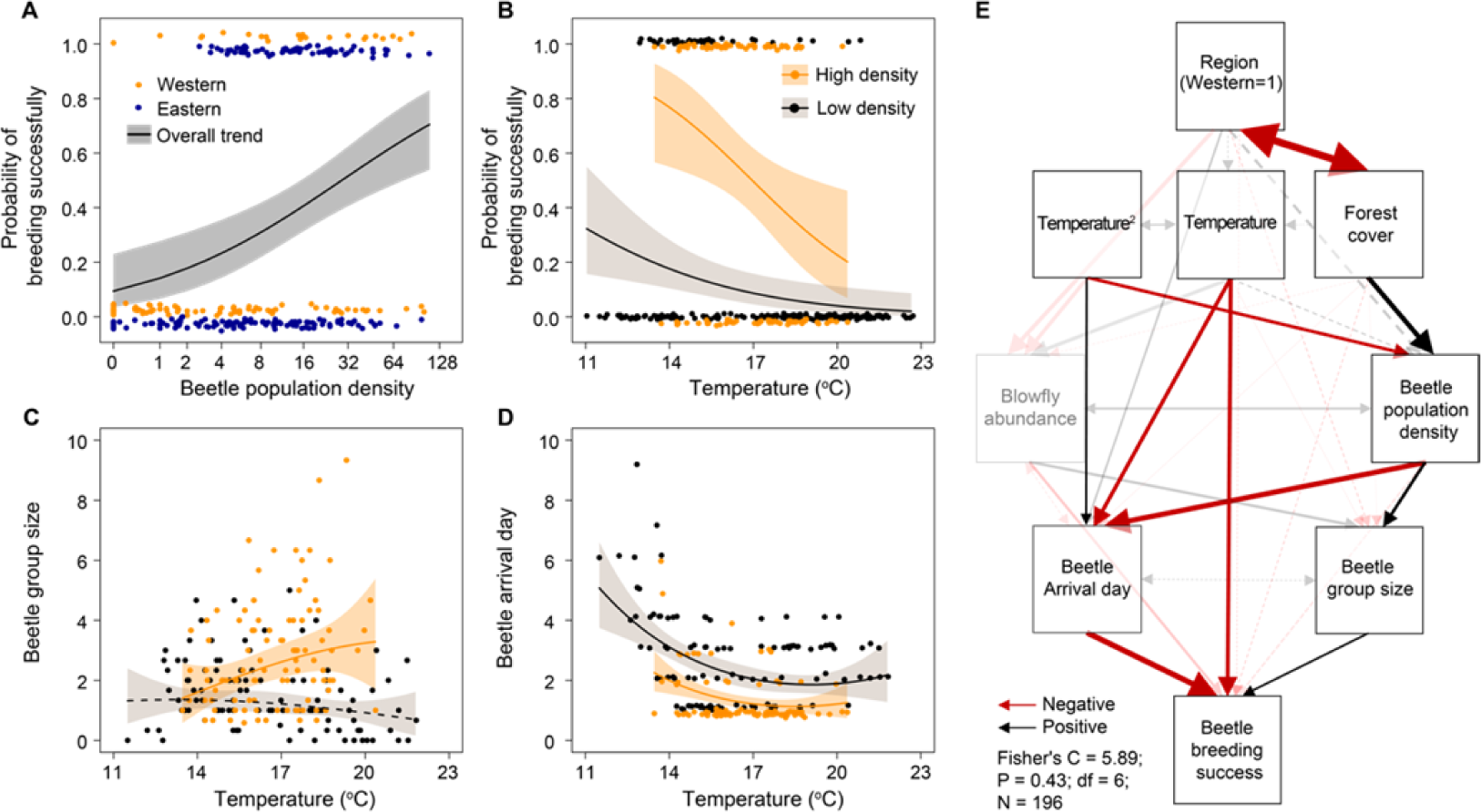
The influence of population density on burying beetle breeding success in field breeding experiments. (A) The influence of population density on beetle breeding success along the western and eastern slopes. (B) The influence of temperature on breeding success at high and low densities. (C) The influence of temperature on group size at high and low densities. (D) The influence of temperature on the arrival day of beetles, representing the day the beetles arrived at the carcass, at high and low densities. (E) Piecewise structural equation modeling (pSEM) for factors affecting breeding success of beetles. In B-D, we considered a cutoff between high and low density at 11 individuals (the rounded mean value of all samples) for a better illustration of the results. All trends and 95% confidence intervals (shaded areas) are estimated from GLMMs. In A, B, & D, points are jittered along the vertical axes to avoid overplotting and improve readability. In E, relationships of interests are highlighted with arrows in black (positive) and dark red (negative), whereas other potential relationships are shown with semitransparent arrows. Unidirectional arrows represent causal relationships, and bidirectional arrows represent correlational relationships. Numbers next to the arrows represent standardized coefficients. Arrows with solid lines are significant (p<0.05), but those with dashed lines are not. The thickness of lines is proportional to the associated standardized coefficients. The influences of year and Julian date were statistically controlled in all causal relationships and were not shown in the figure. See table S7 for the summary of the pSEM.

Next, we compared the elevational distribution of burying beetles on the two slopes with different degrees of land-use change. We found that burying beetles on the western slope (with greater land-use change) were less dense at lower elevations than on the eastern slope (with lower land-use change), and that the lower boundary of the western elevation distribution (1531 m, 20.0°C) was much higher than that of the eastern slope (1219 m, 22.8°C) (GLMM, interaction Elevation*Region, P=0.03, n=372; Fig. 5; Table S3B & S8). However, the upper boundary of the burying beetle distribution was similar on the two slopes (western: 2802 m, 12.5°C; eastern: 2873 m, 11.8°C) (Fig. 5). That is, the thermal niche space, in terms of beetle distribution, on the western slope (with greater land-use change) was much narrower than that on the eastern slope (with lower land-use change), and was is pushed into cooler niche space.

**Figure 5.**
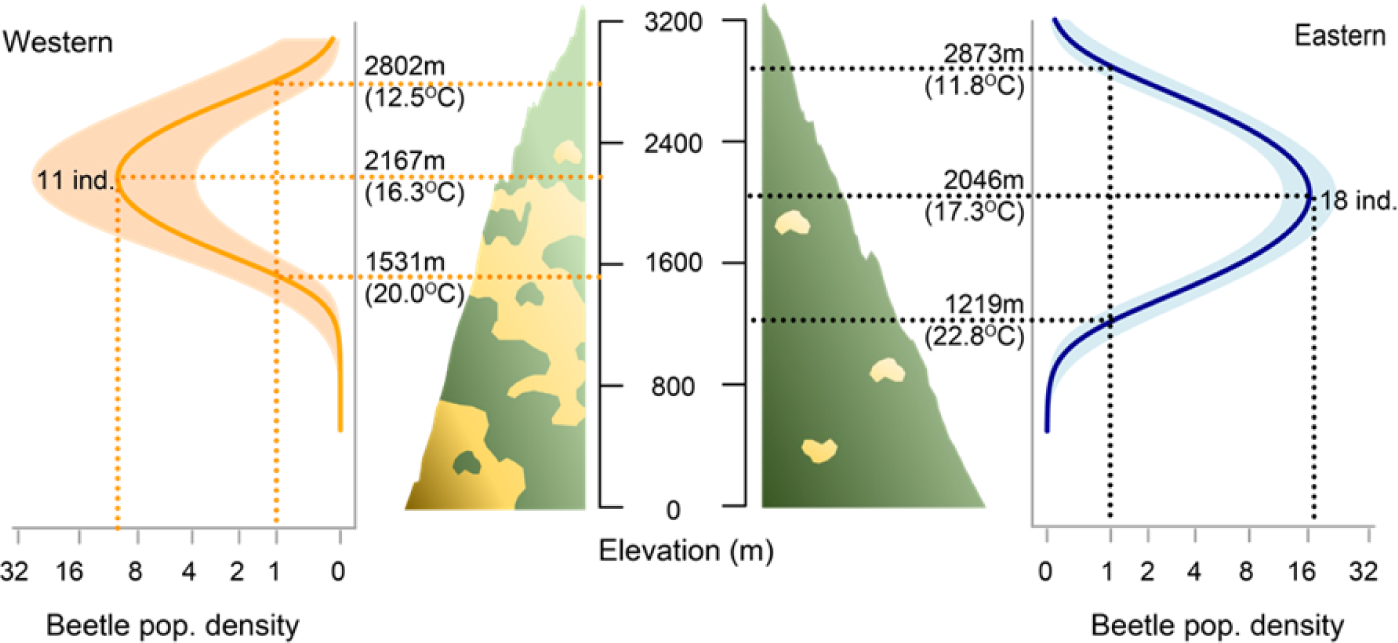
Difference in distributions and thermal niches of burying beetles along the two slopes. Trends and 95% confidence intervals (shaded areas) are estimated from GLMMs. For each slope, the upper elevational limit, elevation with the highest population density, and the lower elevational limit are indicated by dashed lines. The upper and lower limits are the highest and lowest elevations at which the estimated population density (by GLMM) is no less than one individual.

Finally, we experimentally manipulated burying beetle group sizes to simulate the effect of changing beetle population densities in order to determine if population density impacts the beetles’ breeding thermal niche. We found that the probability of breeding successfully for larger groups (simulating group size at high population density) peaked at lower elevations (western: 2244m; eastern: 2247m) and higher temperatures (western: 15.74°C; eastern: 15.24°C), whereas the probability of breeding successfully for smaller groups (simulating group size at low population density) peaked at higher elevations (western: 2757m; eastern: 2628m) and lower temperatures (western: 14.34°C; eastern: 13.64°C; Figure 6A). We also found that on both slopes, larger groups had higher reproductive success at low elevations (GLMM, interaction Elevation*Region*Treatment in the full model, P=0.37; interaction Elevation*Treatment in the best model, P<0.01, n=333; Fig. 6A; Table S9) and high temperatures (GLMM, interaction Temperature*Region*Treatment in the full model, P=0.99; interaction Temperature*Treatment in the best model, P<0.01, n=333; Fig. 6B & C; Table S10) because intraspecific cooperation enables them to outcompete their competitors, blowflies (Chen *et al*. 2020; Liu *et al*. 2020). Experimentally lowering beetle group size on the eastern slope reduced beetle reproductive success, a result that simulates the situation in a fragmented forest, but increasing beetle group size on the western slope recovered their reproductive success from the decline in human-altered habitats. These experimental results further suggest that land-use change can alter the burying beetles’ thermal niche via the Allee effect.

**Figure 6.**
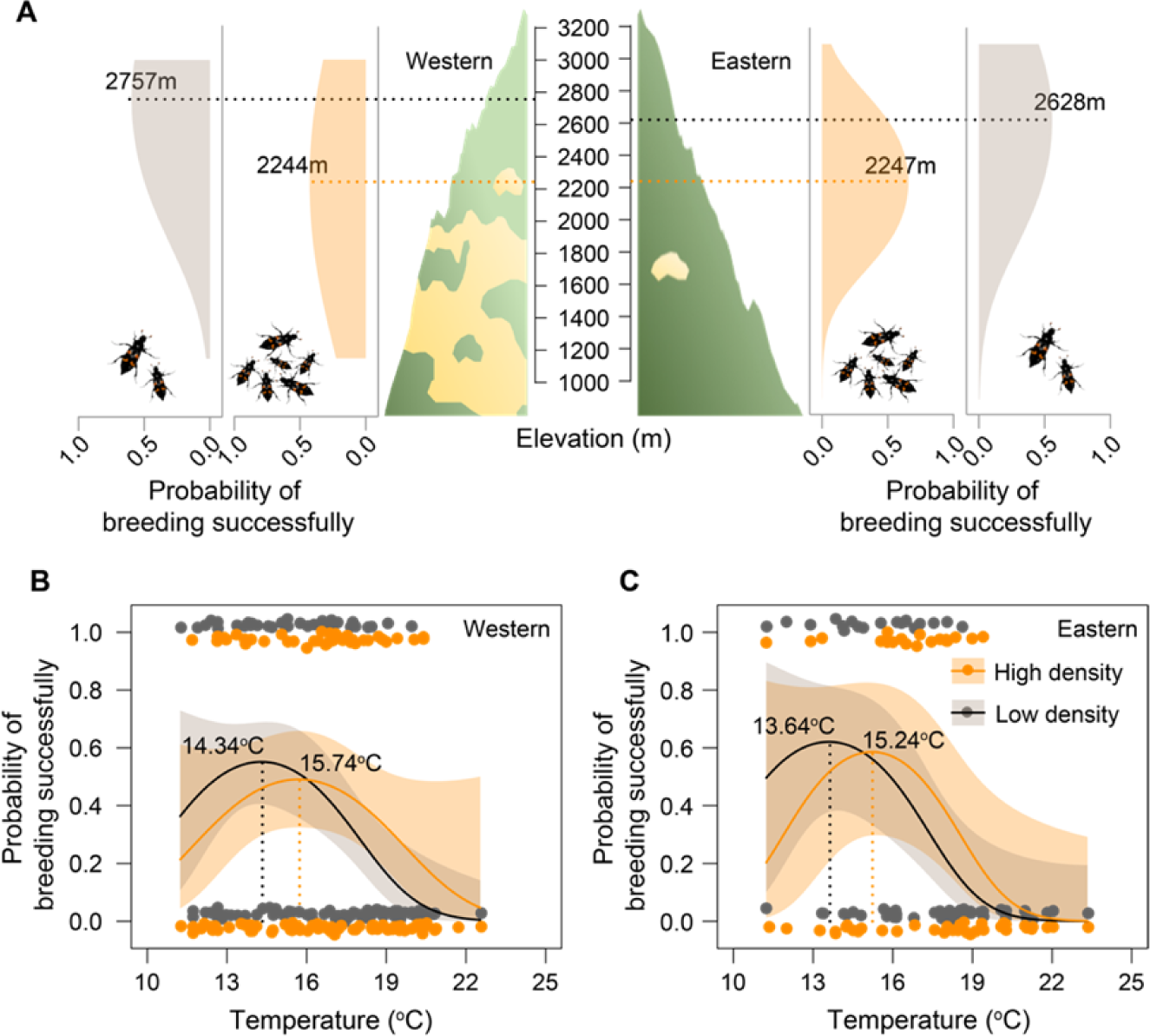
Field manipulative experiments for burying beetle population density. (A) Differences in elevational trends in reproductive success of beetles between treatments (high and low densities) and slopes (western and eastern). (B) The difference in thermal performances of beetles between treatments on the western slope. (C) The difference in thermal performances of beetles between treatments on the eastern slope. In a, the probability of breeding successfully was estimated by GLMM. Each horizontal dashed line indicates the elevation with the highest breeding success for each treatment and region. In B, & C, all trends and confidence intervals (shaded areas) are estimated from GLMM. Vertical dashed lines indicate optimal temperatures, and points were jittered along the vertical axes to avoid overplotting and improve readability.

## Discussion

Our combined theoretical and empirical results suggest that the relationship between biotope and niche is not just a reciprocal correspondence between multidimensional niche space and the physical space in which species live. Instead, we showed that when habitat (biotope) availability decreases, the population density of the organism also decreases, which through the Allee effect (e.g., in our case, the inability of burying beetles to form cooperative groups to compete against blowflies), leads to a reduction in niche width, which in turn leads to their extirpation at lower elevations. To the best of our knowledge, the concept of the Allee effect has not been applied to studying a species’ thermal niche—in this case, the temperature-dependent reproductive success. Thus, this is the first study to experimentally show that changing habitat availability affects a species’ niche width.

Our results indicate that land-use change (i.e., forest loss and fragmentation) has led to a decrease in the population density of *N. nepalensis*, resulting in lower optimal breeding temperatures (thermal niche). Therefore, the reproductive success of *N. nepalensis* in the remaining pristine forests on the slope with greater land-use change is significantly lower than on the slope that maintains large areas of pristine forest cover, even if the microclimatic conditions are the same on both slopes. This has led to the extirpation of *N. nepalensis* on the highly exploited slope, where the lower limit of the elevational distribution of *N. nepalensis* is more than 300 m higher than on the slope with intact forests. Our study further demonstrates that the behavioral mechanisms responsible for the poorer performance of low-density *N. nepalensis* in high-temperature environments include the inability to quickly find reproductive resources (i.e., carcasses) and to form large, cooperative groups for successful reproduction, two behaviors reported in previous studies that enable *N. nepalensis* to overcome extreme ecological and competitive challenges (Sun *et al*. 2014; Chan *et al*. 2019; Chen *et al*. 2020; Liu *et al*. 2020; Tsai *et al*. 2020a). Our study thus provides clear experimental evidence of how land-use change increases the vulnerability of *N. nepalensis* to climate change by reducing its thermal niche through an Allee effect rather than simply by reducing habitat availability and connectivity.

In addition, our study also shows that decreasing population density can cause a reduction in the niche space of species through the Allee effect. For example, our study reveals that under low population density, *N. nepalensis* in warmer areas can no longer cooperate to fight against interspecific competitors. Such an inability to outcompete blowflies reduces the beetles’ reproductive success, and therefore *N. nepalensis* are at greater risk of local extinction in low elevation habitats. Although the Allee effect has been widely applied to understand various behavioral, ecological, and conservation issues (Courchamp *et al*. 1999; Stephens & Sutherland 1999; Angulo *et al*. 2007; Angulo *et al*. 2018), it is worth noting that the Allee effect can be applied not only to social organisms, but also to other species as long as population density can have a positive effect on population growth rate. For example, high population densities make it easier to find mates (Régnière *et al*. 2013), and even trees in a forest can intercept water from fog more efficiently than those in isolation (Dawson 1998); both can be considered examples of the Allee effect. Although an increasing number of theoretical studies have begun to investigate the effects of positive interactions between conspecific individuals on ecological niches (Holt 2009; Koffel *et al*. 2021), empirical studies testing the Allee effect on species niche size remain scarce.

Based on our results, we believe that preserving sufficient areas of suitable habitat to restore population densities to levels sufficient to enhance their breeding performance will help organisms to utilize the remaining suitable habitat to avoid going locally extinction (Soule 1986; Fagan & Holmes 2006). Ultimately, our study shows that anthropogenetic land-use changes may exacerbate the ecological impacts of climate change through the Allee effect. Therefore, understanding the behavioral mechanisms that shape the relationship between habitat availability and thermal niches will be critical to developing effective conservation strategies in this era of climate change.

## Materials and methods

### Individual-based model

To model the effect of land-use changes on population density and thermal niche of social organisms, we set a spatial range of 400 patches and randomly selected the suitable forest habitats according to the habitat ratio. For continuous forest habitats, the habitat ratio is set to 1, which means all 400 patches are forests. For fragmented habitats, the habitat ratio is set to 0.7, which means that 280 patches were randomly selected as forest habitats, and the rest of the patches are non-forested (Fig. 1a). For each simulation, the starting population size was set to 0.6 times the number of patches (i.e., 240 individuals) and was randomly distributed throughout the space.

At each time step of the simulation, the breeding resources (i.e., carcasses) presented randomly in 70% of the forest patches. We then modeled the following three biological processes: (i) reproduction, (ii) dispersal, and (iii) mortality. Notably, in suitable habitats with a carcass, cooperating individuals in the same patch turned the carcass into group resources (i.e., manage and bury the carcass) and reproduced. However, in unsuitable habitats or suitable habitats without carcasses, individuals simply died and could not reproduce, and therefore reproduction was skipped in the simulation. For simplicity, we consider asexual reproduction.

To model the reproduction, we first modeled the effect of environmental temperature on the performance of a solitary individual in a non-cooperative situation by the thermal performance curve (TPC):

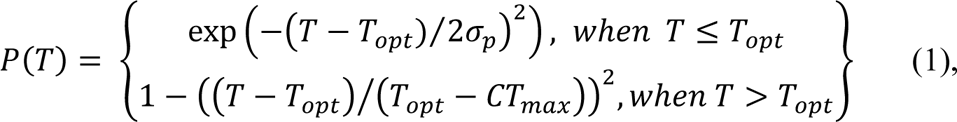

where T is the environment temperature, *σ_p_* is the shape parameter determining the steepness of the curve at the lower end, T_opt_ is the optimum temperature of the species, and CT_max_ is the critical thermal maximum of the species (Deutsch *et al*. 2008; Vasseur *et al*. 2014).

Next, we determined the overall increases in work efficiency due to cooperation

(*E_c_*) by:

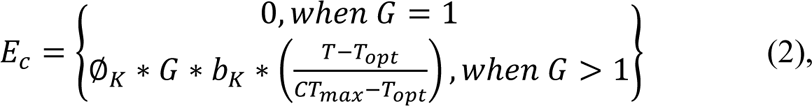

where Ø*_K_* denotes the proportion of energy an individual invests in cooperation, *G* denotes group size (number of individuals in each patch), and *b_K_* denotes an increase in the working efficiency of an individual by cooperation. Notably, for nonsocial species, Ø*_K_ = 0* and hence Ø*_e_ = 0*.

Then, we determined the reproductive increment rate by cooperation (*I*) by a saturating function of *E_e_*:

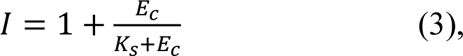

where *K_s_* is the half-saturation constant, which is the value of *E_e_* at which the reproductive increment rate by cooperation (*I*) is half of its maximum. For nonsocial species, *I* = 1 since *E_e_ = 0*, which indicates no increment by cooperation.

Finally, we obtained the productivity of an individual at a given environmental temperature (*T*) by:

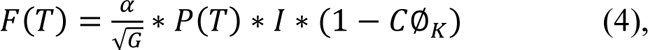

where *α* is the maximum productivity of a solitary individual, and *C* denotes the rate of cost of cooperation. Specifical in nonsocial species, productivity is solely governed by the TPC and the negative density-dependent regulation (i.e., the larger the group size, the lower the productivity) since Ø*_K_* = 0 and *I* = 1.

After an individual produced offspring, the offspring randomly dispersed into patches, which is modeled by a Gaussian kernel density function. Finally, we determined the survival rate (*r*) of the individuals after dispersal by an exponential decay function:

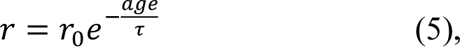

where *r_O_* denotes initial survival rate, *age* refers to the number of survival steps of an individual at the given time step, and *τ* is the exponential time constant that determines the decay rate of survival rates across ages.

The total number of time steps in the simulation is 2,000 to ensure that the system reaches a steady state. We repeated the simulation 200 times for each environmental condition and calculated the mean and standard deviation of the 200 results. Definitions and values for all parameters in our model are summarized in Table S11.

### Field surveys to estimate beetle population densities

We surveyed the population densities of burying beetles by setting hanging pitfall traps (n=372) along the two elevational gradients: 500 to 3275 m above sea level (hereafter, “a.s.l.”) on the western slope (n=148) and 124 to 3209 m a.s.l. on the eastern slope (n=224), with elevation intervals less than 100 m. We conducted these surveys from June to September 2014 to 2015. It is to be noted that these survey samples overlap with those used in previous studies (Chan *et al*. 2019; Liu *et al*. 2020). The hanging pitfall trap consists of a plastic bottle with 100g of rotten meat bait, a landing platform for beetles, and a roof to prevent rainfall (Chan *et al*. 2019; Liu *et al*. 2020; Tsai *et al*. 2020b). We hung each trap in a tree at 1 to 1.5 m above the ground, retrieved it after four nights, and calculated the number of burying beetles as our measure of population density.

### Field breeding experiments

We conducted breeding experiments to test the breeding performance of the beetles in pristine forests (n=392) along the two elevational gradients: 1166 to 2903 m a.s.l. on the western slope (n=176) and 1173 to 2838 m a.s.l. on the eastern slope (n=216), with elevation intervals less than 100m. We conducted these experiments from June to September 2012 to 2015. These experimental samples also partly overlap with those used in previous studies (Chan *et al*. 2019; Liu *et al*. 2020; Tsai *et al*. 2020a). However, we took additional measurements from the video records and different environmental variables for the purposes of this study. Each experimental setup consists of a plastic pot filled with soil, covered by a 2 x 2 cm wire mesh cage to exclude mammalian scavengers (Chan *et al*. 2019; Liu *et al*. 2020; Tsai *et al*. 2020a). We provided a 75g rat carcass as bait (i.e., breeding resource) in the soil-filled pot. Such a design allows burying beetles and blowflies in the environment to freely access the carcass and reproduce. We visited each experiment daily until the carcass was buried by the beetles, decomposed by microbial activity, or consumed by blowfly maggots and other insects (Chan *et al*. 2019). Once the carcass was buried by beetles, we checked the experiment after another 14 days to determine the beetles’ breeding success (i.e., the occurrence of third-instar larvae). We also monitored each experiment with a surveillance camera and a digital video recorder (DVR) to record the burying beetle arrival day (i.e., the day of the first arrival; was successfully identified in 258 of the 259 replicates that the beetles appeared on the carcasses), group size (i.e., the average number of beetles at 22:00, 01:00, and 04:00 on the first night that beetles appeared on the carcass (Liu *et al*. 2020) ; was successfully recorded in 217 of the 259 replicates that the beetles appeared on the carcasses), as well as blowfly abundance on the carcass (i.e., the average number of blowflies counted per two hours on the first day (Chan *et al*. 2019; Chen *et al*. 2020); was successfully recorded in 317 of the 392 replicates). To test the influence of beetle population density on breeding performance, we also surveyed the population density at each site before and after each breeding experiment was carried out in 2014 and 2015 (n=323). We estimated the population density at the beginning of each breeding experiment through linear interpolation by the time between the densities observed before and after the experiment. To prevent mutual interference, successive surveys and experiments in the same location were temporally separated with a minimum of a four-day interval.

### Breeding experiments to examine the influence of local population density

We manipulated breeding group size to reflect the effect of population density on the reproductive performance of burying beetles since higher population densities often result in larger breeding groups under natural conditions. We can thus test whether high population density can rescue the high breeding failure at low elevations on the western slope. We carried out the experiments in pristine forests (n=333) along the two elevational gradients: 1164 to 3000 m a.s.l. on the western slope (n=215) and 900 to 3100 m a.s.l. on the east slope (n=118). These experimental samples also overlap with those used in previous studies (Sun *et al*. 2014; Liu *et al*. 2020; Tsai *et al*. 2020a). Our experimental device comprised an inner plastic container (21 × 13 × 13 cm, containing 10 cm depth of soil) placed within an outer plastic container (41 × 31 × 21.5 cm container, containing 11 cm depth of soil). We provided several entrances on the side wall of the inner container, which allow beetles to freely move between the two containers. The cap of the box was lifted 10 cm above the upper edge of the box to create entrances for blowflies. However, these entrances were also covered by iron mesh (2 × 2 cm cell sizes) to exclude mammalian scavengers. We set up extended eaves around the edge of the cap of the outer container to prevent wild beetles from flying into the container. We also smeared Vaseline on the extended eaves and the side wall of the outer container to prevent wild beetles from climbing into the container. This device allows us to manipulate group size in the container by releasing a specific number of beetles into the container, without being interfered with by wild beetles.

For each set of the experiments, we set up two devices as described above, each containing a 75g rat carcass for beetles. We then released beetles into the containers. One contained one male and one female (i.e., the low-density treatment; n=168), and the other contained three males and three females (i.e., the high-density treatment; n=165). The difference in sample size between the two treatments is because one failed (mainly due to mammalian scavenger invasion) in specific pairs of experiments, resulting in 19 unpaired experiments (12 high-density treatments and seven low-density treatments). We captured the beetles for each set of experiments from nearby locations using the same pitfall traps for our density surveys. However, at low elevations on the western slope, where there was no beetle population, we used beetles captured from the lowest elevation that beetles appeared. To mimic arrival patterns at different elevations, we released the beetles into the experimental device 1, 2, and 3 days after the beginning of each experiment (i.e., placing the rat carcass) at elevations of 1700–2000 m (low), 2000– 2400 m (intermediate) and 2400–2800 m (high), respectively (Sun *et al*. 2014; Liu *et al*. 2020; Tsai *et al*. 2020a). We also visited each experiment every day, following the same protocols as the natural breeding experiments described above.

### Forest cover

We obtained data for forest cover from published global maps of forest cover with a 30-m spatial resolution (Hansen *et al*. 2013). We created a buffer with a 1000 m radius around each sampling point (for both surveys and experiments; n=71) and calculated the mean forest cover within the buffer.

### Environmental temperature monitoring

We recorded the ambient temperature every 30 min for each survey and experiment using an iButton data logger (Maxim Integrated Products, Sunnyvale, CA, USA), which was hung at 1 to 1.5-meters high near the experimental device. We obtained mean daily temperature by averaging the recorded temperature throughout each experiment.

### Data analysis

Depending on the error structure of the dependent variables (DVs), we used Linear Mixed Models (LMM) and Generalized Linear Mixed Models (GLMM) in the R package *lme4* (Bates *et al*. 2015) to analyze the relationships among variables. We carried out LMMs for continuous DVs (e.g., temperature, forest cover, and blowfly abundance), binomial GLMMs for binary DVs (e.g., presence and breeding success of beetles), and negative binomial GLMM for count data (e.g., population density). Some continuous DVs were transformed for normality prior to analysis: blowfly abundance was log-transformed, and forest cover was square root transformed. All independent variables (IVs) were standardized to facilitate model convergence. Julian date was considered as a covariate (control variable) in all models. For all analyses, we simultaneously included the main effects of all IVs. However, for complicated models with interactions among IVs or quadric terms (namely for testing the nonlinear elevation or temperature), we began with a full model, but sequentially removed non-significant (P>0.05) interactions and quadric terms to obtain a parsimonious model. Since many of our sites were sampled more than once, we considered “sampling site” as a random factor to control for repeated sampling. However, for the manipulative breeding experiments that were carried out in matched pairs, we assigned each pair of experiments a unique serial number and used this serial number as a random factor nested within sampling site (coded as 1|sampling site/serial number, in the language of R package *lme4*). To control for potential among-year variation, we also considered year as a crossed random factor for those analyses involving data obtained from more than three years, but considered year as a fixed factor for these analyses involving data obtained from only two years. We obtained statistical significances from type II sums of squares for models without interactions, but from type III sums of squares models with interactions, all of which were carried out using the R package *car* (Fox & Weisberg 2011). We also calculated *R^2^* for each model via the method described by Nakagawa & Schielzeth (Nakagawa & Schielzeth 2013) using the R package *MuMIn* (Barton 2020).

We also conducted piecewise structure equation modeling (pSEM) (Shipley 2009; Lefcheck 2016) to examine the potential causal relations among variables. The analyses involved three major steps: (i) creating a hypothetical causal framework; (ii) testing the goodness-of-fit of the hypothetical framework by d-sep tests (Shipley 2009); and (iii) testing the relationships among variables under the hypothetical framework. We also controlled for the influences of year and Julian date for the potential causal relations. For the parts of pSEMs involving beetle population density as a dependent variable, we used log-transformed density and LMM to perform the analysis in order to obtain standardized coefficients for the potential causal relationships. For any relationship involving a binary response (i.e., breeding success), the standardized coefficient was estimated manually following the approach proposed by Menard (Menard 2004).

Specifically for the results of natural breeding experiments, sample sizes vary among analyses due to misfunction in the temperature monitoring or video recording equipment, or the lack of corresponding population density surveys, in some of these experiments. We describe these sample size variations individually in supplementary tables for the results of analyses.

### Research Permits

Permits for the field study were issued by Taiwan’s Forestry Bureau (1014107656, 1024103748, 1034102881, and 1044105071 by Nantou Forest District Office; 1018106554, 1028101879, and 1038102225 by Hualien Forest District Office; 1013106573, 1023240523, 1033102695, and 1043103919 by Tungshih Forest District Office), and Taroko National Parks (201207310237, 201304120275, 201404090315, 201506020389) from 2012 to 2015.

### Data availability

The C++ based program for the individual-based model is available in Zenodo (10.5281/zenodo.7039827). All data generated in this study are deposited on Dryad (Chan, Shih-Fan et al. (2022), Allee effects mediate the impact of land-use change on the thermal niche of social species, Dryad, Dataset, https://doi.org/10.5061/dryad.9s4mw6mkz). In addition, R code for data analysis (https://doi.org/10.5281/zenodo.7181290) and supplementary information (https://doi.org/10.5281/zenodo.7181292) are available in Zenodo.

## Acknowledgments

We thank staff at the Mei-Feng Highland Experiment Farm and Taroko National Park for the logistical support for field experiments. We also thank Yen-Cheng Lin, Tzu-Neng Yuan, Ching-Fu Lin, Bo-Fei Chen, Yu-Ching Liu, and Mark Liu for their support in the field. SFS was supported by Academia Sinica (AS-SS-106-05) and Minister of Science and Technology of Taiwan (100-2621-B-001-004-MY3, 104-2311-B-001-028-MY3, and 108-2314-B-001-009-MY3). DRR was supported by the National Science Foundation (IOS-1656098).

## Supplementary Information

### Supplementary tables

**Table S1.** Regression tables of the liner model for differences in forest cover along elevational gradients between the eastern and western slopes.

|  | Estimate | SE | Z | P |
| --- | --- | --- | --- | --- |
| Intercept | 1.27 | 0.02 | 68.03 | <0.001 |
| Region (western=1) | -0.32 | 0.03 | -10.63 | <0.001 |
| Elevation | -0.12 | 0.03 | -4.00 | <0.001 |
| Elevation<sup>2</sup> | -0.38 | 0.05 | -7.96 | <0.001 |
| Region*Elevation | 0.23 | 0.05 | 5.05 | <0.001 |
| Region*Elevation<sup>2</sup> | 0.46 | 0.09 | 5.35 | <0.001 |

$R^2=0.72$ ; N=71

**Table S2.** Regression tables of LMM for differences in environmental temperatures along elevational gradients between the eastern and western slopes.

|  | Estimate | SE | t | P |
| --- | --- | --- | --- | --- |
| Intercept | 16.86 | 0.08 | 213.43 | <0.001 |
| Region (western=1) | -0.12 | 0.11 | -1.071 | 0.29 |
| Elevation | -3.19 | 0.08 | -41.85 | <0.001 |
| Region*Elevation | -0.19 | 0.11 | -1.74 | 0.09 |
| Year | -0.39 | 0.02 | -21.48 | <0.001 |
| Julian Date | -0.38 | 0.02 | -23.61 | <0.001 |
| Julian Date<sup>2</sup> | -0.36 | 0.01 | -25.91 | <0.001 |

$R^2=0.91$ ; n=4849 observations from 37 plots

**Table S3.** Regression table of GLMM for the effects of forest cover, temperature, and regions on bury beetle population densities. (A) Testing for the joint influences of temperature and forest cover. (B) Testing for the difference in the effects of temperature between the easter and western slopes.

|  | Estimate | SE | Z | P |
| --- | --- | --- | --- | --- |
| Intercept | 1.89 | 0.19 | 10.05 | <0.001 |
| Temperature | -0.17 | 0.14 | -1.26 | 0.21 |
| Temperature<sup>2</sup> | -1.03 | 0.13 | -8.19 | <0.001 |
| Forest cover | 0.69 | 0.16 | 4.29 | <0.001 |
| Temperature*Forest cover | 0.44 | 0.14 | 3.10 | <0.01 |
| Year | -0.30 | 0.07 | -4.27 | <0.001 |
| Julian date | -0.26 | 0.06 | -4.41 | <0.001 |

$R^2=0.65$ ; n=372 B
|  | Estimate | SE | Z | P |
| --- | --- | --- | --- | --- |
| Intercept | 1.87 | 0.20 | 9.17 | <0.001 |
| Temperature | -0.08 | 0.14 | -0.55 | 0.58 |
| Temperature<sup>2</sup> | -0.97 | 0.12 | -8.13 | <0.001 |
| Region (western=1) | -0.50 | 0.17 | -2.90 | <0.01 |
| Temperature*Region | -0.25 | 0.15 | -1.70 | 0.09 |
| Year | -0.30 | 0.07 | -4.05 | <0.001 |
| Julian date | -0.28 | 0.06 | -4.67 | <0.001 |

$R^2=0.50$ ; n=372

**Table S4.** Regression tables of GLMM for the influence of beetle population density on the beetles’ breeding success.

|  | Estimate | SE | Z | P |
| --- | --- | --- | --- | --- |
| Intercept | -1.03 | 0.28 | -3.75 | <0.001 |
| Population density | 1.72 | 0.44 | 3.93 | <0.001 |
| Temperature | -2.19 | 0.49 | -4.46 | <0.001 |
| Temperature<sup>2</sup> | -1.99 | 1.14 | -1.76 | 0.08 |
| Population density*Temperature | -2.08 | 0.99 | -2.11 | 0.04 |
| Julian date | -0.51 | 0.34 | -1.51 | 0.13 |
| Year | 0.86 | 0.34 | -2.51 | 0.01 |

$R^2=0.47$ ; n=322\*
\*We excluded one of the 333 replicates because of missing temperature data due to malfunction in the equipment.

**Table S5.** Regression table of LMM for the influence of beetle population density on burying beetles’ group size.

|  | Estimate | SE | t | P |
| --- | --- | --- | --- | --- |
| Intercept | 1.26 | 0.08 | 16.44 | <0.001 |
| Population density | 0.29 | 0.08 | 3.46 | <0.001 |
| Temperature | -0.19 | 0.09 | -2.10 | 0.04 |
| Temperature<sup>2</sup> | -0.37 | 0.15 | -2.42 | 0.02 |
| Population density*Temperature | 0.56 | 0.17 | 3.23 | <0.01 |
| Julian date | -0.32 | 0.08 | -4.18 | <0.001 |
| Year | 0.06 | 0.10 | 0.63 | 0.53 |

$R^2=0.30$ ; n=199\*
\*We excluded one of the 217 replicates because of missing temperature data due to malfunction in the equipment, and 17 of the remaining replicates conducted before 2014 since population density was not surveyed.

**Table S6.** Regression table of LMM for the influence of beetle population density on burying beetles’ arrival day on the carcasses.

|  | Estimate | SE | t | P |
| --- | --- | --- | --- | --- |
| Intercept | 1.27 | 0.03 | 39.71 | <0.001 |
| Population density | -0.41 | 0.06 | -7.05 | <0.001 |
| Temperature | -0.25 | 0.05 | -4.71 | <0.001 |
| Temperature<sup>2</sup> | 0.32 | 0.09 | 3.40 | <0.001 |
| Population density*Temperature | 0.17 | 0.09 | 1.86 | 0.07 |
| Julian date | -0.05 | 0.05 | -0.99 | 0.32 |
| Year | -0.18 | 0.05 | -3.64 | <0.001 |

$R^2=0.48$ ; n=210\*
\*We excluded seven of the 258 replicates because of missing temperature data due to misfunction in the equipment, and 41 of the remaining replicates conducted before 2014 since population density was not surveyed.

**Table S7.** Results of piecewise SEM for factors affecting the bury beetles’ breeding success.

| Relationship between variables |  | Estimate | Standardized Estimate | SE | P |
| --- | --- | --- | --- | --- | --- |
| **Causal relations:** |  |  |  |  |  |
| Beetle breeding success<br>($R^2=0.43$) | ← Region | -0.31 | -0.05 | 0.52 | 0.55 |
|  | ← Forest cover | -0.44 | -0.07 | 0.49 | 0.38 |
|  | ← Beetle arrival day | -2.95 | -0.47 | 0.53 | <0.001 |
|  | ← Beetle group size | 1.21 | 0.19 | 0.46 | <0.01 |
|  | ← Temperature | -2.05 | -0.33 | 0.48 | <0.001 |
|  | ← Blowfly abundance | -1.43 | -0.23 | 0.51 | <0.01 |
|  | ← Julian date | -0.60 | -0.10 | 0.43 | 0.17 |
|  | ← Year | 0.04 | 0.01 | 0.45 | 0.93 |
| Beetle group size<br>($R^2=0.32$) | ← Region | -0.12 | -0.12 | 0.09 | 0.20 |
|  | ← Forest cover | -0.06 | -0.06 | 0.09 | 0.52 |
|  | ← Beetle pop. density | 0.34 | 0.34 | 0.08 | <0.001 |
|  | ← Blowfly abundance | 0.19 | 0.19 | 0.08 | 0.02 |
|  | ← Temperature | -0.10 | -0.10 | 0.07 | 0.14 |
|  | ← Julian date | -0.23 | -0.23 | 0.07 | <0.001 |
|  | ← Year | 0.11 | 0.11 | 0.07 | 0.125 |
| Beetle arrival day<br>($R^2=0.52$) | ← Region | 0.19 | 0.19 | 0.07 | <0.01 |
|  | ← Forest cover | -0.02 | -0.02 | 0.07 | 0.82 |
|  | ← Beetle pop. density | -0.42 | -0.42 | 0.07 | <0.001 |
|  | ← Temperature | -0.38 | -0.38 | 0.05 | <0.001 |
|  | ← Temperature<sup>2</sup> | 0.37 | 0.22 | 0.11 | 0.001 |
|  | ← Julian date | -0.07 | -0.07 | 0.06 | 0.24 |
|  | ← Year | -0.15 | -0.15 | 0.06 | 0.02 |
| Beetle population density<br>($R^2=0.28$) | ← Region | 0.25 | 0.25 | 0.14 | 0.09 |
|  | ← Forest cover | 0.42 | 0.42 | 0.14 | <0.01 |
|  | ← Temperature | 0.11 | 0.11 | 0.07 | 0.13 |
|  | ← Temperature<sup>2</sup> | -0.48 | -0.28 | 0.10 | <0.001 |
|  | ← Julian date | -0.23 | -0.23 | 0.05 | <0.001 |
|  | ← Year | -0.03 | -0.03 | 0.06 | 0.62 |
| Blowfly abundance<br>($R^2=0.40$) | ← Region | -0.34 | -0.34 | 0.09 | <0.001 |
|  | ← Forest cover | 0.07 | 0.07 | 0.08 | 0.40 |
|  | ← Temperature | 0.32 | 0.32 | 0.06 | <0.001 |
|  | ← Temperature<sup>2</sup> | -0.58 | -0.34 | 0.11 | <0.001 |
|  | ← Julian date | -0.25 | -0.25 | 0.06 | <0.001 |
|  | ← Year | -0.30 | -0.30 | 0.07 | <0.001 |
| Temperature<br>($R^2=0.03$) | ← Region | -0.06 | -0.06 | 0.22 | 0.78 |
|  | ← Forest cover | -0.04 | -0.04 | 0.22 | 0.87 |
|  | ← Julian date | -0.14 | -0.14 | 0.04 | <0.001 |
|  | ← Year | -0.11 | -0.11 | 0.05 | 0.03 |
| **Correlations:** |  |  |  |  |  |
| Forest | ↔ Region | -0.65 | -0.65 | - | <0.001 |
| Temperature<sup>2</sup> | ↔ Temperature | 0.16 | 0.16 | - | 0.01 |
| Blowfly abundance | ↔ Beetle pop. density | 0.22 | 0.22 | - | <0.001 |
| Beetle arrival day | ↔ Beetle group size | 0.02 | 0.02 | - | 0.37 |
| Beetle arrival day | ↔ Blowfly abundance | -0.08 | -0.08 | - | 0.15 |

Fisher's C = 5.98; P-value = 0.43; df=6; n=196\*
\* We included those replicates without any missing data in the analysis.

**Table S8.**
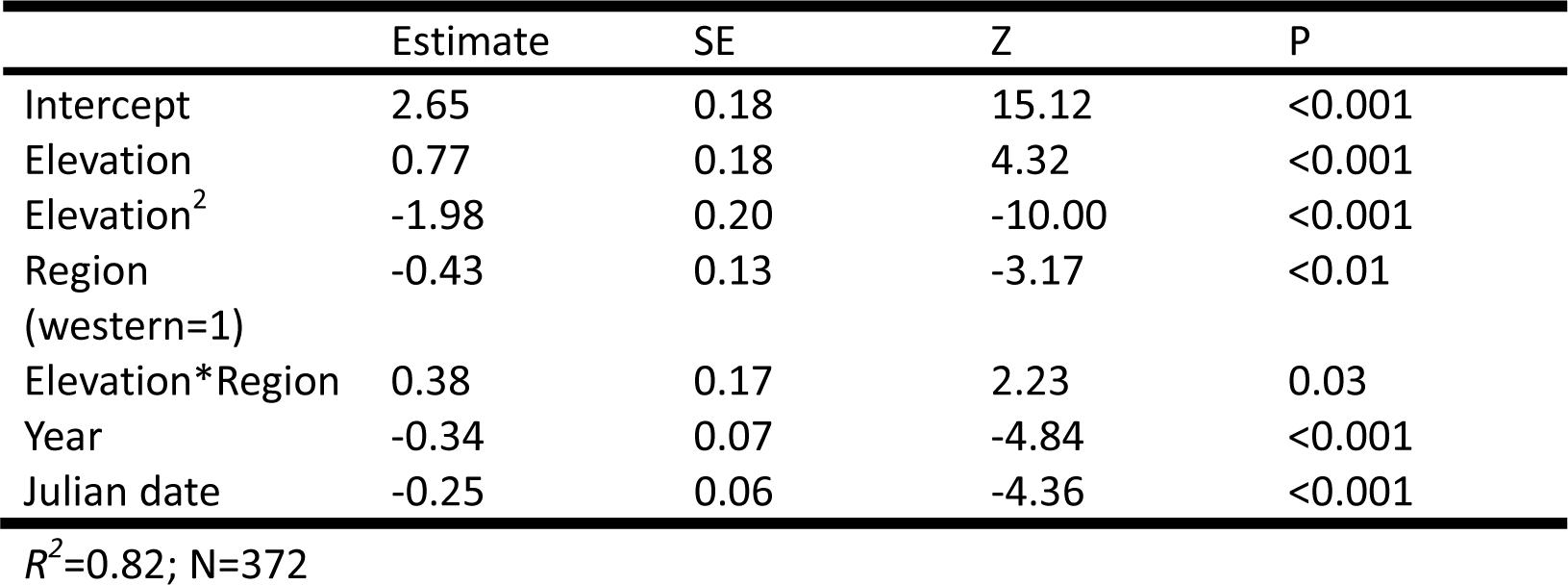
Regression table of GLMM for testing the between-slope difference in the beetles’ elevational distribution.

**Table S9.**
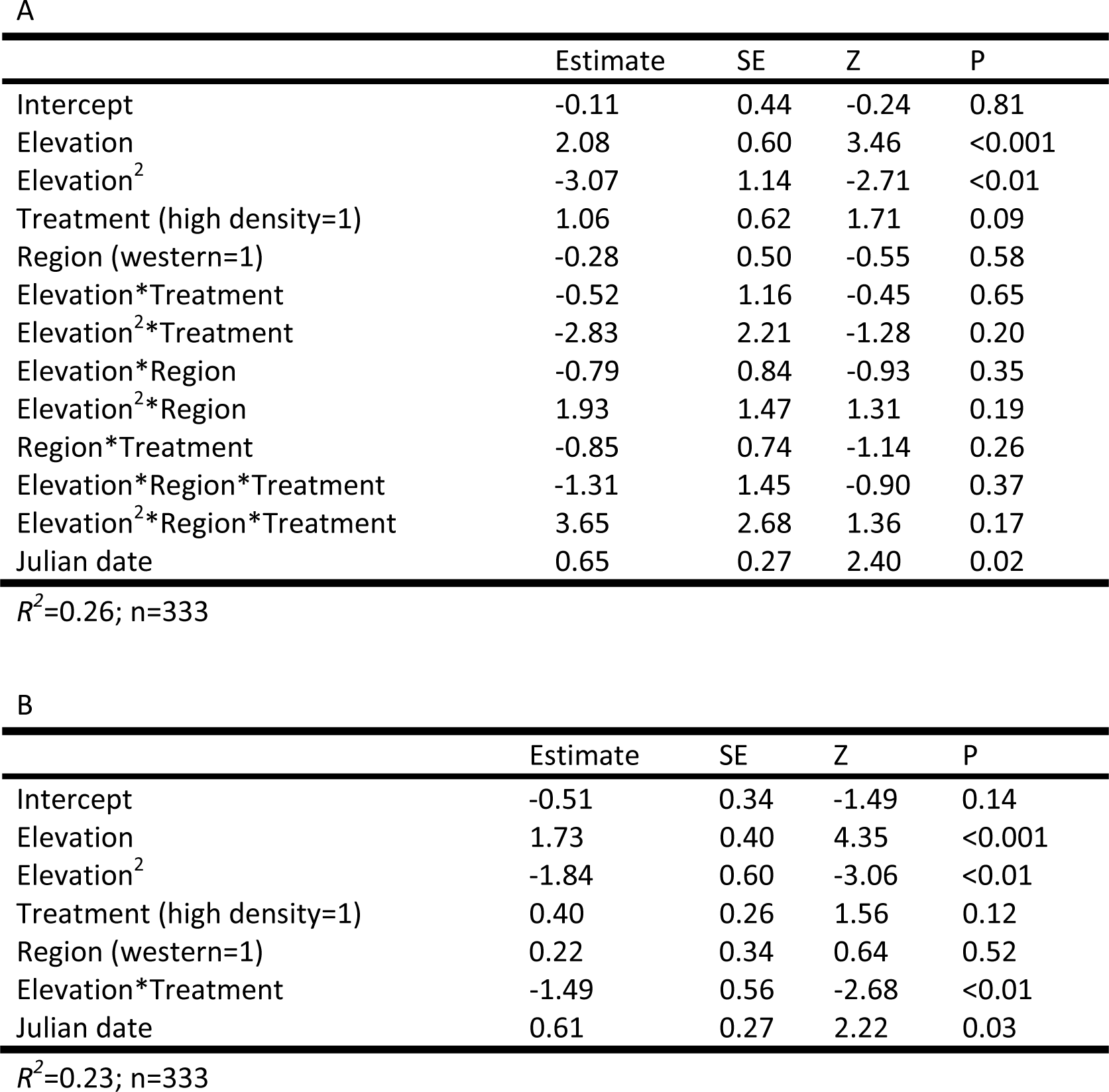
Regression table of GLMM for factors affecting elevational trends in breeding success in manipulative experiments. (A) A full model considering the interaction among treatment, elevation, and region. (B) Reduced model retaining only significant interactions.

|  | Estimate | SE | Z | P |
| --- | --- | --- | --- | --- |
| Intercept | -0.11 | 0.44 | -0.24 | 0.81 |
| Elevation | 2.08 | 0.60 | 3.46 | <0.001 |
| Elevation<sup>2</sup> | -3.07 | 1.14 | -2.71 | <0.01 |
| Treatment (high density=1) | 1.06 | 0.62 | 1.71 | 0.09 |
| Region (western=1) | -0.28 | 0.50 | -0.55 | 0.58 |
| Elevation*Treatment | -0.52 | 1.16 | -0.45 | 0.65 |
| Elevation<sup>2</sup>*Treatment | -2.83 | 2.21 | -1.28 | 0.20 |
| Elevation*Region | -0.79 | 0.84 | -0.93 | 0.35 |
| Elevation<sup>2</sup>*Region | 1.93 | 1.47 | 1.31 | 0.19 |
| Region*Treatment | -0.85 | 0.74 | -1.14 | 0.26 |
| Elevation*Region*Treatment | -1.31 | 1.45 | -0.90 | 0.37 |
| Elevation<sup>2</sup>*Region*Treatment | 3.65 | 2.68 | 1.36 | 0.17 |
| Julian date | 0.65 | 0.27 | 2.40 | 0.02 |

$R^2=0.26$ ; n=333 B
|  | Estimate | SE | Z | P |
| --- | --- | --- | --- | --- |
| Intercept | -0.51 | 0.34 | -1.49 | 0.14 |
| Elevation | 1.73 | 0.40 | 4.35 | <0.001 |
| Elevation<sup>2</sup> | -1.84 | 0.60 | -3.06 | <0.01 |
| Treatment (high density=1) | 0.40 | 0.26 | 1.56 | 0.12 |
| Region (western=1) | 0.22 | 0.34 | 0.64 | 0.52 |
| Elevation*Treatment | -1.49 | 0.56 | -2.68 | <0.01 |
| Julian date | 0.61 | 0.27 | 2.22 | 0.03 |

$R^2=0.23$ ; n=333

**Table S10.** Regression table of GLMM for factors affecting thermal trends in breeding success in manipulative experiments. (A) A full model considering the interaction among treatment, mean daily temperature, and region. (B) Reduced model retaining only significant interactions.

|  | Estimate | SE | Z | P |
| --- | --- | --- | --- | --- |
| Intercept | -0.26 | 0.29 | -0.89 | 0.37 |
| Temperature | -2.24 | 0.70 | -3.20 | <0.01 |
| Temperature<sup>2</sup> | -2.92 | 1.15 | -2.54 | 0.01 |
| Treatment (High density=1) | 0.82 | 0.56 | 1.48 | 0.14 |
| Region (western=1) | 0.07 | 0.35 | 0.19 | 0.85 |
| Temperature*Treatment | 1.63 | 1.38 | 1.19 | 0.24 |
| Temperature<sup>2</sup>*Treatment | -1.12 | 2.30 | -0.49 | 0.63 |
| Temperature*Region | 0.75 | 0.82 | 0.92 | 0.36 |
| Temperature<sup>2</sup>*Region | 0.67 | 1.38 | 0.49 | 0.63 |
| Region*Treatment | -0.59 | 0.69 | -0.86 | 0.39 |
| Temperature*Region*Treatment | 0.02 | 1.62 | 0.011 | 0.99 |
| Temperature<sup>2</sup>*Region*Treatment | 1.93 | 2.76 | 0.70 | 0.49 |
| Julian date | 0.18 | 0.27 | 0.65 | 0.51 |

$R^2 = 0.32$ ; n=333 B
|  | Estimate | SE | Z | P |
| --- | --- | --- | --- | --- |
| Intercept | -0.28 | 0.25 | -1.11 | 0.27 |
| Temperature | -1.71 | 0.37 | -4.70 | <0.001 |
| Temperature<sup>2</sup> | -2.46 | 0.64 | -3.82 | <0.001 |
| Treatment (high density=1) | 0.45 | 0.26 | 1.71 | 0.09 |
| Region (western=1) | 0.10 | 0.27 | 0.35 | 0.73 |
| Temperature*Treatment | 1.55 | 0.59 | 2.62 | <0.01 |
| Julian date | 0.16 | 0.27 | 0.59 | 0.56 |

$R^2 = 0.28$ ; n=333

**Table S11.** Summary of parameters in the individual-based model.

| Name | Value | Description |
| --- | --- | --- |
| $T$ | [17, 23] | Environmental temperature |
| $\sigma_p$ | 2.0 | Shape parameter determining the steepness of the curve at the lower end |
| $T_{opt}$ | 16.7 | Optimal temperature in TPC |
| $CT_{max}$ | 23.0 | Critical thermal maximum in TPC |
| $\phi_K$ | 0.5 | Degree of cooperation |
| $b_K$ | 10 | Increase in efficiency by cooperation |
| $K_s$ | 2 | Half-saturation constant |
| $\alpha$ | 2 | Maximum productivity of an individual |
| $C$ | 0.1 | Rate of cost of cooperation |

### Supplementary figures

**Figure 1— Figure Supplement 1.**
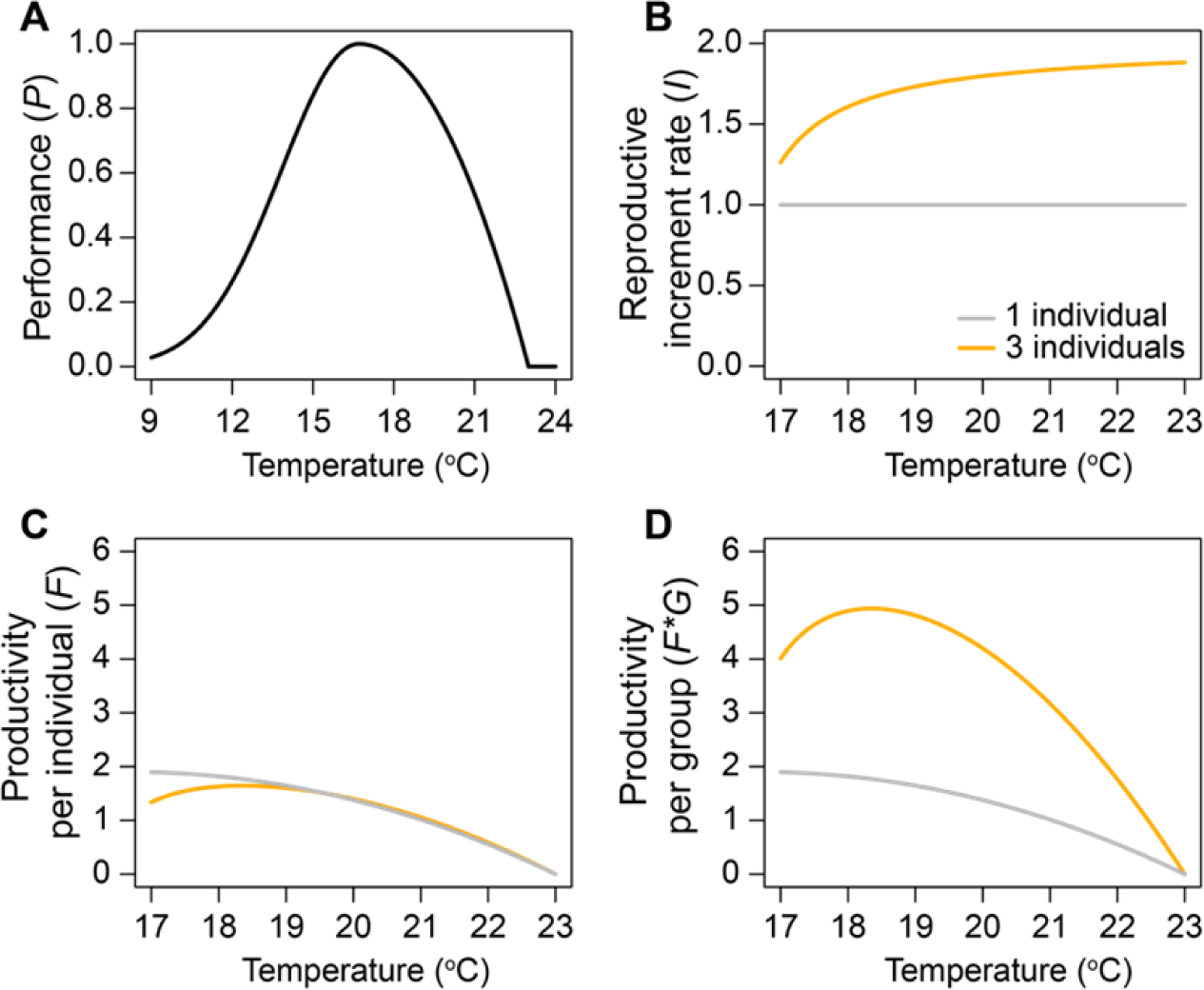
Settings for individual-based models. (A) The thermal performance curve used in the model. (B) Variation in cooperative benefits along the temperature gradient. (C) Variation in individual productivity in small and large groups along the temperature gradient. (D) Variation in group productivity in small and large groups along the temperature gradient.

**Figure 1— Figure Supplement 2.**
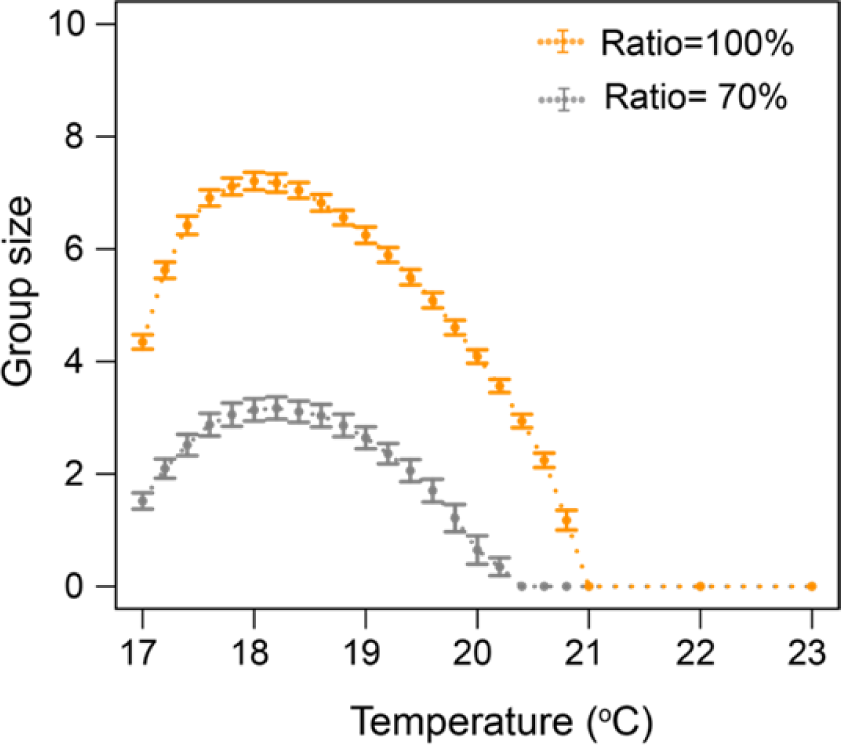
Simulation results from the individual-based model for the impacts of land-use change on group sizes of social populations. Points and error bars represent means and standard deviations obtained from 200 simulations.

**Figure 1— Figure Supplement 3.**
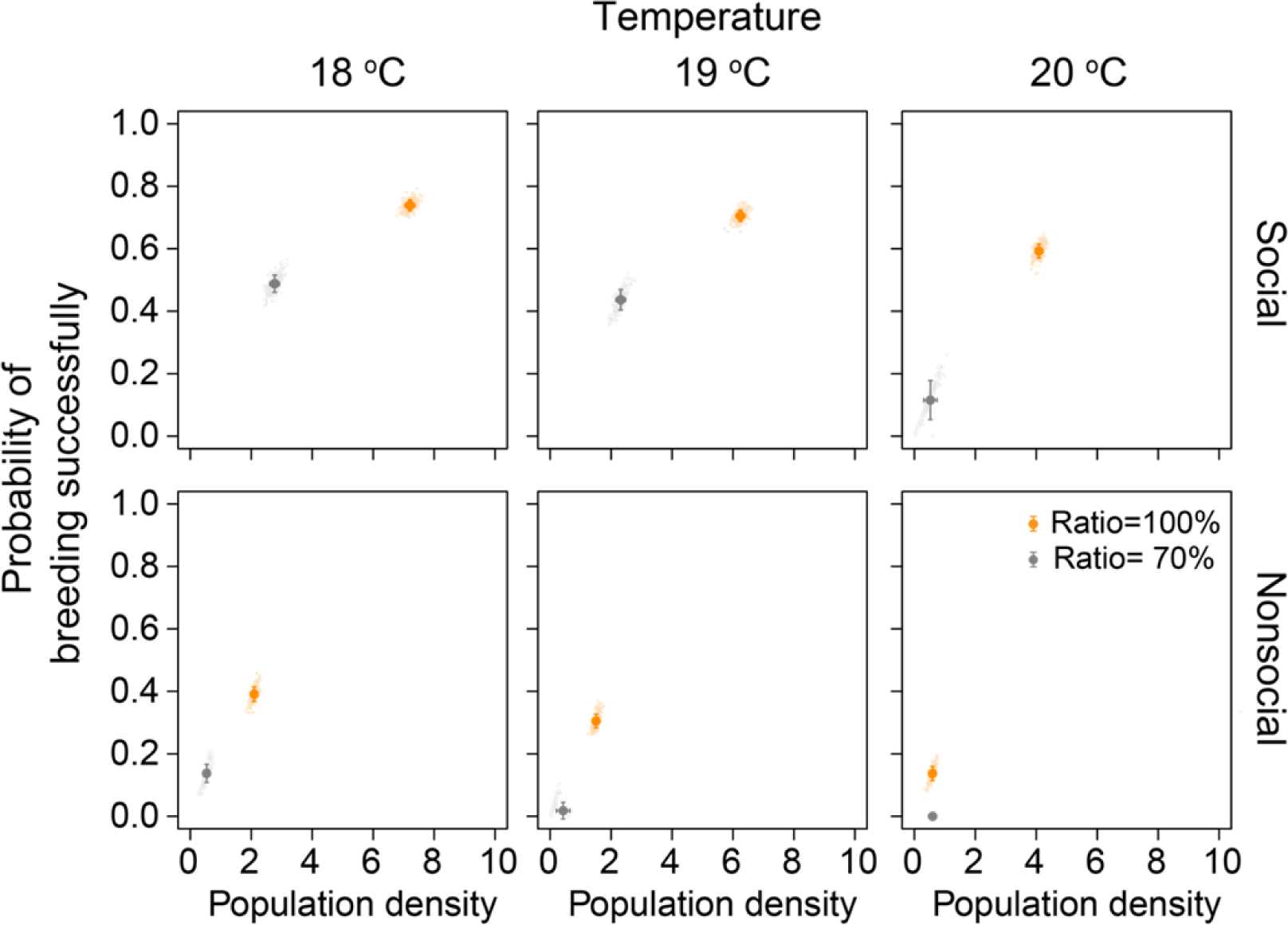
Simulation results from the individual-based model for the effects of population density on the probability of breeding successfully. Points represent outcomes from the simulations. Solid circles and error bars represent means and standard deviations.

